# preon: Fast and accurate entity normalization for drug names and cancer types in precision oncology

**DOI:** 10.1101/2023.05.22.540912

**Authors:** Arik Ermshaus, Michael Piechotta, Gina Rüter, Ulrich Keilholz, Ulf Leser, Manuela Benary

## Abstract

**Motivation:** In precision oncology, clinicians are aiming to find the best treatment for any patient based on their molecular characterization. A major bottleneck is the annotation and evaluation of individual variants, for which usually a range of knowledge bases are manually screened. To incorporate and integrate the vast information of different databases, fast and accurate methods for harmonization are necessary.

**Summary:** preon is a fast and accurate library for the normalization of drug names and cancer types in large-scale data integration.

**Availability and Implementation:** preon is implemented in Python and freely available via the PyPI repository. Source code and gold standard data sets are available at https://github.com/ermshaua/preon/.

**Supplementary information:** Supplementary data are available online.

---

Precision oncology (PO) considers the molecular make-up of cancer patients for therapy decisions and promises better-targeted therapies. It requires extensive knowledge bases about associations of molecular features, cancer types, and drugs, which are typically created by integrating multiple specialized databases to leverage international community efforts [6, 3, 7]. Such an integration requires the accurate normalization of biomedical entities from different databases into a common ontology [5]. As databases may contain tens of thousands of such entities, a normalization algorithm must be fast and accurate. In contrast to entity normalization in texts [1], it can only rely on syntactic name features, as the to-be-normalized entities are extracted from database columns instead of natural language sentences.

We present preon, a Python-library for drug name and cancer type normalization in data integration projects. To balance speed and accuracy, preon is based on an efficient multi-step process performing a cascade of matching algorithms of increasing complexity (see Figure 1A). As preprocessing, preon transforms a given entity name by extracting its alphanumeric characters and applies a lowercase transformation. As a first matching step, preon tries to match the reduced sequence exactly to its pre-processed reference dictionary. If not successful, preon next performs token or n-gram based name matching to allow for different subword orders and slight variations. If still unsuccessful, preon calculates the pairwise edit distances to all reference names. Steps two and three apply individual thresholds to define matches. As reference sets, preon uses ChEMBL [2] for drug names and the cancer subtree in disease ontology [4] for tumor entities.

**Fig. 1.**
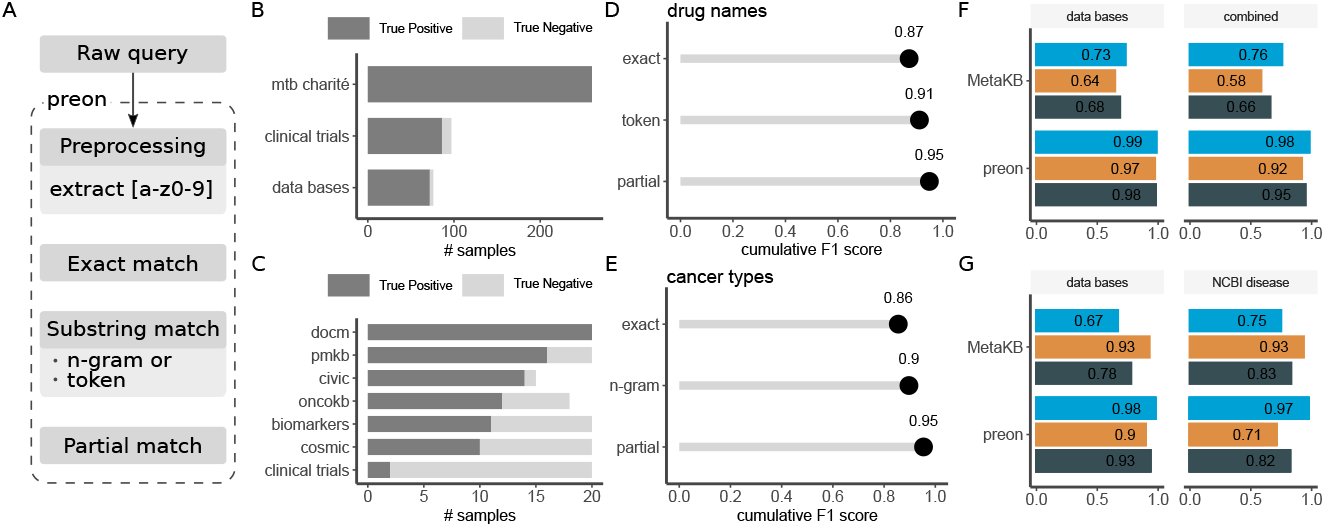
**A** The term normalization workflow established in preon. The normalization consists of a multi-step matching process. Overview of the gold standard for **B** drug names and **C** cancer types. The increase in F1 score when using exact, token, and partial matching is depicted in **D** for drug name normalization (combined data set) and in **E** for cancer types. **F** and **G** performance comparison using preon and the normalization presented in MetaKB [7] for drug names and cancer types, respectively. Precision (blue), recall (yellow), and F1-score (dark gray) are shown for different gold standards (see supp. material for details).

## Evaluation

For the evaluation, we developed gold standard sets for drug names as well as cancer types. The former is based on clinical reports and different databases (see Figure 1B) while the latter uses the NCBI Disease corpus and different databases (see Figure 1C). Details of the gold standards are available in supporting material (Table S1 and S2).

We performed a precision-recall analysis and calculated the combined F1-score to evaluate the effects of the three different matching steps in preon. We also evaluated runtimes and compared results to the term normalization performed in MetaKB [7], a large integration project for precision oncology. MetaKB performs normalization using mainly web queries with BioThings [9] and ChEMBL [2].

For drug name normalization, exact matching yields an F1-score of 87%. N-gram matching improves the F1-score by 4pp (percent points) and edit-distance-based matching adds another 4pp (see Figure 1D). The best pre-defined similarity threshold for partial matching was established at 20% in our gold standard. In cancer type normalization, exact matching reaches an F1-score of 86%, improved by 4pp through n-gram matching and further 5pp through edit-distance-based matching (Figure 1E) also using a threshold of 20%. preon outperforms MetaKB on all measures for drug name normalization (see Figure 1F), with an increase in F1 of almost 30pp. For cancer type normalization, preon has higher precision yet lower recall than MetaKB for both gold standards (see Figure 1G). In terms of F1, preon outperforms MetaKB by 15pp in the database data set and is on-par (−1pp) in the NCBI disease data set.

preon also is very fast because the costly steps two and especially three are only applied when the previous steps found no match, which very often is not the case. preon requires 53ms on average for normalizing a single drug name and 7ms for cancer types in the evaluation (see supp. Table S3). When applied to full databases, preon requires, for instance, 14s / 7s to normalize all 2.8k drug names / 3.5k cancer types from CIVIC. In contrast, such a normalization with MetaKB is not feasible as it is implemented as a web service.

## Conclusion

preon is an accurate library for normalizing drug names and cancer types and it is fast enough to be applied in large PO integration projects. Critical decisions for building and assessing such a library are the choice of reference library and the construction of the gold standard for evaluation. Regarding the former, also DrugBank for drug names and MeSH for cancer types would be viable candidates; actually, we provide access to both databases in our implementation and examples how to easily exchange the reference set in preon. Regarding the latter, our four gold standard data sets have the limitation that the fractions of true negatives to true positives are somewhat arbitrary, although they largely influence accuracy and runtime measurements, especially because entity names without a match are particularly expensive to - not - normalize. We thus believe that an international effort to create appropriate evaluation data would be an important step for the future. Furthermore, we believe that preon’s runtime can be improved further by using advanced indexing techniques for the second and third step [8].

## References

1. Kim, D., Lee, J., So, C.H., Jeon, H., Jeong, M., Choi, Y., Yoon, W., Sung, M., Kang, J.: A neural named entity recognition and multi-type normalization tool for biomedical text mining. IEEE Access 7, 73729–73740 (2019)

2. Mendez, D., Gaulton, A., Bento, A.P., Chambers, J., Veij, M.D., Félix, E., Magariños, M.P., Mosquera, J.F., Mutowo, P., Nowotka, M., Gordillo-Marañón, M., Hunter, F., Junco, L., Mugumbate, G., Rodriguez-Lopez, M., Atkinson, F., Bosc, N., Radoux, C.J., Segura-Cabrera, A., Hersey, A., Leach, A.R.: ChEMBL: Towards direct deposition of bioassay data. Nucleic Acids Research 47, D930–D940 (1 2019). https://doi.org/10.1093/nar/gky1075,http://www.pathogenbox.org

3. Pallarz, S., Benary, M., Lamping, M., Rieke, D., Starlinger, J., Sers, C., Wiegandt, D.L., Seibert, M., Ševa, J., Schäfer, R., Keilholz, U., Leser, U.: Comparative analysis of public knowledge bases for precision oncology. JCO Precision Oncology pp. 1–8 (11 2019). https://doi.org/10.1200/PO.18.00371, http://ascopubs.org/doi/10.1200/PO.18.00371

4. Schriml, L.M., Mitraka, E., Munro, J., Tauber, B., Schor, M., Nickle, L., Felix, V., Jeng, L., Bearer, C., Lichenstein, R., Bisordi, K., Campion, N., Hyman, B., Kurland, D., Oates, C.P., Kibbey, S., Sreekumar, P., Le, C., Giglio, M., Greene, C.: Human Disease Ontology 2018 update: Classification, content and workflow expansion. Nucleic Acids Research 47, D955–D962 (1 2019). https://doi.org/10.1093/nar/gky1032, https://pubmed.ncbi.nlm.nih.gov/30407550/

5. Sharp, M.E.: Toward a comprehensive drug ontology: Extraction of drug-indication relations from diverse information sources. Journal of Biomedical Semantics 8 (1 2017). https://doi.org/10.1186/s13326-016-0110-0

6. Starlinger, J., Pallarz, S., Ševa, J., Rieke, D., Sers, C., Keilholz, U., Leser, U.: Variant information systems for precision oncology. BMC Medical Informatics and Decision Making 18, 107 (11 2018). https://doi.org/10.1186/s12911-018-0665-z, https://bmcmedinformdecismak.biomedcentral.com/articles/10.1186/s12911-018-0665-z

7. Wagner, A.H., Walsh, B., Mayfield, G., Tamborero, D., Sonkin, D., Krysiak, K., Deu-Pons, J., Duren, R.P., Gao, J., McMurry, J., Patterson, S., del Vecchio Fitz, C., Pitel, B.A., Sezerman, O.U., Ellrott, K., Warner, J.L., Rieke, D.T., Aittokallio, T., Cerami, E., Ritter, D.I., Schriml, L.M., Freimuth, R.R., Haendel, M., Raca, G., Madhavan, S., Baudis, M., Beckmann, J.S., Dienstmann, R., Chakravarty, D., Li, X.S., Mockus, S., Elemento, O., Schultz, N., Lopez-Bigas, N., Lawler, M., Goecks, J., Griffith, M., Griffith, O.L., Margolin, A.A.: A harmonized meta-knowledgebase of clinical interpretations of somatic genomic variants in cancer. Nature Genetics 52, 448–457 (4 2020). https://doi.org/10.1038/s41588-020-0603-8, https://doi.org/10.1038/s41588-020-0603-8

8. Wandelt, S., Deng, D., Gerdjikov, S., Mishra, S., Mitankin, P., Patil, M., Siragusa, E., Tiskin, A., Wang, W., Wang, J., Leser, U.: State-of-the-art in string similarity search and join. SIGMOD Record 43(1), 64–76 (2014)

9. Xin, J., Afrasiabi, C., Lelong, S., Adesara, J., Tsueng, G., Su, A.I., Wu, C.: Cross-linking BioThings APIs through JSON-LD to facilitate knowledge exploration. BMC Bioinformatics 19, 30 (2 2018). https://doi.org/10.1186/s12859-018-2041-5, https://bmcbioinformatics.biomedcentral.com/articles/10.1186/s12859-018-2041-5

