## Supplemental Material for "preon: Fast and accurate entity normalization for drug names and cancer types in precision oncology"

### Supplemental material and methods

May 17, 2023

#### Contents

**1 Description of gold standard for drug name normalization** **1**
1.1 Process of manual annotation and file description . . . . . . . . . 2

**2 Description of gold standard for cancer type normalization** **2**
2.1 Process of manual annotation and file description . . . . . . . . . 3

**3 Definition of precision and recall for fuzzy matching** **4**

**4 Identifying thresholds for fuzzy matching** **4**

**5 Dealing with class imbalance** **5**

**6 preon and MetaKB runtimes** **6**

#### 1 Description of gold standard for drug name normalization

As there is no gold standard for drug names and the possibilities of synonyms are extensive, we generated our own gold standard using commonly used names from three different types of sources (see supp. Table S1). First, we used drug names presented in the molecular tumor board at the Charité Comprehensive Cancer Center. Second, we sampled drug names from the data bases Biomarkers, CIViC, oncoKB and TARGET. And last, we used the semi-structured entries from https://clinicaltrials.gov/ in the column "Intervention" to sample another cohort. All common names were matched to the corresponding ChEMBL ID to generate the goldstandard (available at https://github.com/ermshaua/preon/).

These data sets comprise a variety of problems, which will occur in reports especially setting:

1. Spelling issues,
2. Use of synonyms,

| Source | Number of Entities | Avg. Length | Perc. of Multi-Token Names | Number of matched Chembl-IDs |
|---|---|---|---|---|
| MTB Charité | 260 | 9.26 | 6.54 | 260 |
| Data bases | 76 | 11.25 | 9.21 | 72 |
| Clinical Trials | 97 | 15.34 | 34.02 | 86 |
| Combined | 421 | 10.97 | 13.16 | 406 |

Table S1: Number of samples for the gold standard of drug names. The different sources are separately listed.

3. Suggestion of combination therapies,
4. Use of abbreviations, and
5. Use of drug classes rather than a specific medication.

From the original data, we removed all samples which describe a drug class (e.g. BRAF inhibitor), because this is a different type of task. We included combination therapies with the goal to identify a ChEMBL ID for each individual substance.

##### 1.1 Process of manual annotation and file description

Data description:

The table includes 5 columns:

1. source ... The source from where we sampled the drug names for normalization. This can be a database, the annotation from the molecular tumor board or clinicaltrials.gov.
2. treatment ... the treatment given in clinical trials, which can include more than one drug.
3. drug name ... The common name of the drug as taken from the corresponding source.
4. ChEMBL ID ... ID from ChEMBL https://www.ebi.ac.uk/chembl/
5. NCT ID ... for the treatments and drugs coming from clinicialtrials.gov, the corresponding IDs are given.

#### 2 Description of gold standard for cancer type normalization

Within the project, we are trying to integrate multiple databases to provide a systematic overview of therapeutic options for cancer patients based on single nucleotide variants. Some drugs are only effective in certain types of cancers and it is very valuable to provide the cancer entity if this information is available. To make the cancer types comparable, it is useful to normalize common names to standard names. We have randomly selected a number of common cancer names from each database and prepared them for manual annotation.

For the normalization of cancer entities, we relied on two different types of data sources. First, we sampled 20 entities from different databases, respectively, including clinical trials (see supp. Table S2) and matched them manually with the corresponding entry in disease ontology (Schriml et al., 2019). This data set is available as supp. material and the data format is described in more detail in the following section:

#### 2.1 Process of manual annotation and file description

All entries from the different databases were first matched with precise matching to IDs from disease ontology. Only 58 tumor entity names had a unique mapping, with 65 entities having no mapping or a partial mapping to disease ontology. Unclear annotations were validated by a clinician.

Data description:

The table has 4 main parts:

1. source ... The database from where we sampled the common name of tumor entities.
2. cancer type ... The common name of the tumor entity as taken from the disease description in the corresponding database.
3. DOID ... ID from Disease Ontology https://disease-ontology.org/. There can be no entry, one entry or multiple entries separated by comma.
4. None ... these entries do not constitute a tumor entity in the cancer subtree of disease ontology. This is for clarification on behalf of the annotating clinician.

The data rows consists of three parts, marked with different colors:

1. Green ... Cancer types with exactly one identifier in the cancer subtree of disease ontology (n = 82).
2. Yellow ... Cancer types, with multiple identifiers along one level in the cancer subtree of disease ontology (n = 3).
3. Orange ... Entries, which are not defined as tumor entities according to disease ontologies (n = 48).

Second, we used NCBI Disease, which is a data set with abstracts (Doğan et al., 2014) in which diseases are annotated with MESH/OMIM-IDs. We used mondo (Shefchek et al., 2020) to relate the MESH identifiers with the corresponding ones from disease ontology. Because we are focusing on normalization of tumor entities, we reduced the data set by including only diseases from the cancer related subtree (DOID:162).

| Source | Number of Entities | Avg. Length | Perc. of Multi-Token Names | Number of matched DOIDs |
|---|---|---|---|---|
| Biomarkers | 20 | 13.05 | 45.00 | 11 |
| CIViC | 15 | 22.40 | 80.00 | 14 |
| Clinical Trials | 20 | 26.00 | 95.00 | 2 |
| Cosmic | 20 | 20.10 | 15.00 | 10 |
| DOCM | 20 | 20.75 | 85.00 | 20 |
| oncoKB | 18 | 23.06 | 77.78 | 12 |
| PMKB | 20 | 21.35 | 70.00 | 16 |
| NCBI disease | 158 | 22.22 | 75.76 | 158 |

Table S2: Number of samples for the gold standard of cancer types. The different sources are separately listed.

### 3 Definition of precision and recall for fuzzy matching

In preon, we are focusing on the use case, where we are searching for an identifier in a given nomenclature. Based on our gold standard, we can define a contigency table and thus calculate precision (Equation 1), recall (Equation 2) and the harmonic mean of the two, called $F_1$ score (Equation 3).

$$\begin{aligned} Precision &= \frac{TP}{TP+FP} \\ &= 1 - FDR \end{aligned} \quad (1)$$

$$Recall = \frac{TP}{TP+FN} \quad (2)$$

$$\begin{aligned} F_1 &= \frac{2 \cdot Precision \cdot Recall}{Precision + Recall} \\ &= \frac{2 \cdot TP}{2 \cdot TP + FP + FN} \end{aligned} \quad (3)$$

In a fuzzy matching approach, multiple identifiers can be returned. We label a result as a true positive, if at least one identifier is correctly returned. It is up to the medical inclined user to verify the correct identifier.

### 4 Identifying thresholds for fuzzy matching

In a first step, we identified the best number of elements for partial matching (supp. Figure 1A and D) as well as the number of tokens in the n-gram matching (supp. Figure 1D). Using moderate partial matching thresholds (20%-30%) and bigrams increases recall while maintaining high precision.

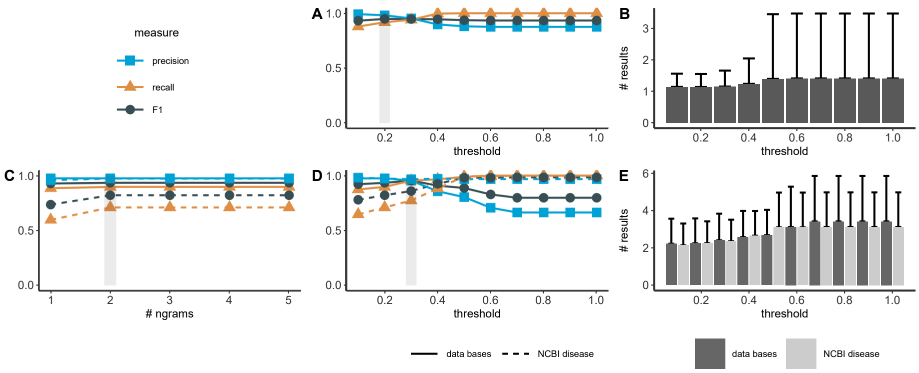


Figure 1: **A** Effect of fuzzy matching for drug names. The threshold is given as a percentage of the length of the drug names. Precision (blue), recall (orange) and F1 sores (dark gray) are shown. The light gray highlights the chosen threshold for further evaluation. **B** Number of results returned for different thresholds in the fuzzy matching. Shown is the mean (bar) and standard deviation as error bar over all drug names. **C** Precision (blue), recall (orange) and F1 scores (dark gray) for different number of n-grams for cancer type normalization. The results for the databases are shown with solid lines, the results for the NCBI disease data set are shown with dashed lines. **D** Effect of fuzzy matching for cancer types similar to **A**. **E** Number of results returned for different thresholds for cancer types similar to **B**

With increasing thresholds for the partial matching, the number of identifiers for the users to check will increase. Thus, we not only optimized the threshold for best precision and recall, but also took the number of results into account (see supp. Figure 1B and E).

### 5 Dealing with class imbalance

The F1-score is dependent on the ratio between positive to negative cases (Williams, 2021). The fraction of positives in the data set can be denoted by

$$\pi = P/(P + N) \tag{4}$$

With $\pi \rightarrow 1$ the precision will increase and converge to 1 (Williams, 2021).

Thus we expanded our gold standard to include more negative cases. For the NCBI data set we have around 300 cancer queries and 1000 queries annotated as true negatives with respect to tumor entities. preon reaches a precision of 76.9%, a recall of 71% and a F1 score of 73.8%. Although precision is lower in this case, it validates the quality of preon and highlights the importance of balanced test data sets. In comparison, MetaKB only has a precision of 9% and a recall of 93%. This dramatic decrease in precision is related to the fact, that MetaKB is not specific for tumor entities but will return any ID from the disease ontology.

### 6 preon and MetaKB runtimes

We measured the runtime for preon and MetaKB for the drug name and cancer type normalisation task. For MetaKB we used https://github.com/cancervariants/metakb. Supp. Table S3 reports the summary statistics for each dataset. On average, preon and MetaKB share similar results for drug name normalisation. However, preon is 15 times faster than MetaKB for cancer type normalisation. preon's differences in runtime between the tasks are mostly explained by the different amount of reference data (around 100k entries in ChEMBL and around 10k entries in DO).

| Dataset | preon | MetaKB |
|---|---|---|
| Drug Combined | (0.05/53/419/22,666) | (19/67/286/28,347) |
| Drug DB | (0.05/38/436/2,91) | (30/73/286/5,567) |
| Cancer DB | (0.07/7/40/971) | (31/107/239/14,011) |
| NCBI Disease | (0.06/9/33/1,48) | (32/128/258/20,218) |

Table S3: Min/Average/Max/Total runtimes in ms for normalisation and matching in preon and MetaKB for queries from the different datasets.

### References


Rezarta Islamaj Doğan, Robert Leaman, et al. NCBI disease corpus: A resource for disease name recognition and concept normalization. *Journal of Biomedical Informatics*, 47:1–10, 2014. ISSN 15320464. doi:10.1016/j.jbi.2013.12.006.

Lynn M. Schriml, Elvira Mitraka, et al. Human Disease Ontology 2018 update: Classification, content and workflow expansion. *Nucleic Acids Research*, 47:D955–D962, 2019. ISSN 13624962. doi:10.1093/nar/gky1032.

Kent A. Shefchek, Nomi L. Harris, et al. The Monarch Initiative in 2019: An integrative data and analytic platform connecting phenotypes to genotypes across species. *Nucleic Acids Research*, 48:D704–D715, 2020. ISSN 13624962. doi:10.1093/nar/gkz997.

Christopher K. I. Williams. The effect of class imbalance on precision-recall curves. *Neural Computation*, 33:853–857, 2021. ISSN 0899-7667. doi:10.1162/neco_a_01362.
